# Rapid selection of a human monoclonal antibody that potently neutralizes SARS-CoV-2 in two animal models

**DOI:** 10.1101/2020.05.13.093088

**Authors:** Wei Li, Aleksandra Drelich, David R. Martinez, Lisa Gralinski, Chuan Chen, Zehua Sun, Alexandra Schäfer, Sarah R. Leist, Xianglei Liu, Doncho Zhelev, Liyong Zhang, Eric C. Peterson, Alex Conard, John W. Mellors, Chien-Te Tseng, Ralph S. Baric, Dimiter S. Dimitrov

**Affiliations:** Center for Antibody Therapeutics, Division of Infectious Diseases, Department of Medicine, University of Pittsburgh Medical School, 3550 Terrace Str, Pittsburgh, PA 15261, USA; Department of Microbiology & Immunology, Centers for Biodefense and Emerging Diseases, Galveston National Laboratory, 301 University Blvd, Galveston, Texas 77550, USA; University of North Carolina at Chapel Hill, 135 Dauer Drive, 3109 Michael Hooker Research Center Chapel Hill, NC 27599; Abound Bio, 1401 Forbes Ave, Pittsburgh, PA 15219

**Keywords:** therapeutic antibodies, coronaviruses, SARS-CoV-2

## Abstract

Effective therapies are urgently needed for the SARS-CoV-2/COVID19 pandemic. We identified panels of fully human monoclonal antibodies (mAbs) from eight large phage-displayed Fab, scFv and VH libraries by panning against the receptor binding domain (RBD) of the SARS-CoV-2 spike (S) glycoprotein. One high affinity mAb, IgG1 ab1, specifically neutralized replication competent SARS-CoV-2 with exceptional potency as measured by two different assays. There was no enhancement of pseudovirus infection in cells expressing Fcγ receptors at any concentration. It competed with human angiotensin-converting enzyme 2 (hACE2) for binding to RBD suggesting a competitive mechanism of virus neutralization. IgG1 ab1 potently neutralized mouse ACE2 adapted SARS-CoV-2 in wild type BALB/c mice and native virus in hACE2 expressing transgenic mice. The ab1 sequence has relatively low number of somatic mutations indicating that ab1-like antibodies could be quickly elicited during natural SARS-CoV-2 infection or by RBD-based vaccines. IgG1 ab1 does not have developability liabilities, and thus has potential for therapy and prophylaxis of SARS-CoV-2 infections. The rapid identification (within 6 days) of potent mAbs shows the value of large antibody libraries for response to public health threats from emerging microbes.

---

The severe acute respiratory distress coronavirus 2 (SARS-CoV-2) (1) has spread worldwide thus requiring safe and effective prevention and therapy. Inactivated serum from convalescent patients inhibited SARS-CoV-2 replication and decreased symptom severity of newly infected patients (2, 3) suggesting that monoclonal antibodies (mAbs) could be even more effective. Human mAbs are typically highly target-specific and relatively non-toxic. By using phage display we have previously identified a number of potent fully human mAbs (m396, m336, m102.4) against emerging viruses including severe acute respiratory syndrome coronavirus (SARS-CoV) (4), Middle East respiratory syndrome coronavirus (MERS-CoV) (5) and henipaviruses (6, 7), respectively, which are also highly effective in animal models of infection (8–11); one of them was administered on a compassionate basis to humans exposed to henipaviruses and successfully evaluated in a clinical trial (12).

Size and diversity of phage-displayed libraries are critical for rapid selection of high affinity antibodies without the need for additional affinity maturation. Our exceptionally potent antibody against the MERS-CoV, m336, was directly selected from very large (size ∼10^11^ clones) library from 50 individuals (5). However, another potent antibody, m102.4, against henipavirusses was additionally affinity matured from its predecessor selected from smaller library (size ∼10^10^ clones) from 10 individuals (7, 13). Thus, to generate high affinity and safe mAbs we used eight very large (size ∼ 10^11^ clones each) naive human antibody libraries in Fab, scFv or VH format using PBMCs from 490 individuals total obtained before the SARS-CoV-2 outbreak. Four of the libraries were based on single human VH domains where CDRs (except CDR1 which was mutagenized or grafted) from our other libraries were grafted as previously described (14).

Another important factor to consider when selecting effective mAbs is the appropriate antigen. Similar to SARS-CoV, SARS-CoV-2 uses the spike glycoprotein (S) to enter into host cells. The S receptor binding domain (RBD) binds to its receptor, the human angiotensin-converting enzyme 2 (hACE2), thus initiating series of events leading to virus entry into cells (15, 16). We have previously characterized the function of the SARS-CoV S glycoprotein and identified its RBD which is stable in isolation (17). The RBD was then used as an antigen to pan phage displayed antibody libraries; we identified potent antibodies (5, 8) more rapidly and the antibodies were more potent than when we used whole S protein or S2 (unpublished). In addition, the SARS-CoV RBD based immunogens are highly immunogenic and elicit neutralizing antibodies which protect against SARS-CoV infections (18). Thus, to identify SARS-CoV-2 mAbs, we generated two variants of the SARS-CoV-2 RBD (aa 330-532) (Fig. S1) and used them as antigens for panning of our eight libraries.

Panels of high-affinity binders to RBD in Fab, scFv and VH domain formats were identified. There was no preferential use of any antibody VH gene (an example for a panel of binders selected from the scFv library is shown in Fig. S2A) and the number of somatic mutations was relatively low (Fig. S2B, for the same panel of binders as in Fig. S2A). For nine of the highest affinity mAbs a provisional patent application was filed on March 12, 2020 by the University of Pittsburgh. Those high affinity mAbs can be divided into two groups in terms of their competition with hACE2. Two representatives of each group are Fab ab1 and VH ab5. To further increase their binding through avidity effects and extend their half-live in vivo they were converted to IgG1 and VH-Fc fusion formats, respectively. Ab1 was characterized in more details because of its potential for prophylaxis and therapy of SARS-CoV-2 infection.

The Fab and IgG1 ab1 bound strongly to the RBD (Fig. 1A) and the whole SARS-CoV-2 S1 protein (Fig. 1B) as measured by ELISA. The Fab ab1 equilibrium dissociation constant, *K*_d_, as measured by the biolayer interferometry technology (BLItz), was 1.5 nM (Fig. 1C). The IgG1 ab1 bound with high (*K*_d_ =160 pM) avidity to recombinant RBD (Fig. 1D). IgG1 ab1 bound cell surface associated native S glycoprotein suggesting that the conformation of its epitope on the RBD in isolation is close to that in the native S protein (Fig. 2, S3). The binding of IgG1 ab1 was of higher avidity than that of hACE2-Fc (Fig. 2B). Binding of ab1 was specific for the SARS-CoV-2 RBD; it did not bind to the SARS-CoV S1 (Fig. 3A) nor to cells that do not express SARS-CoV-2 S glycoprotein (Fig. 2A). Ab1 competed with hACE2 for binding to the RBD (Fig. 3B and C) indicating possible neutralization of the virus by preventing binding to its receptor. It did not compete with the CR3022 (Fig. 3D and E), which also binds to SARS-CoV (19) and with ab5 (Fig. 3F).

**Figure 1.**
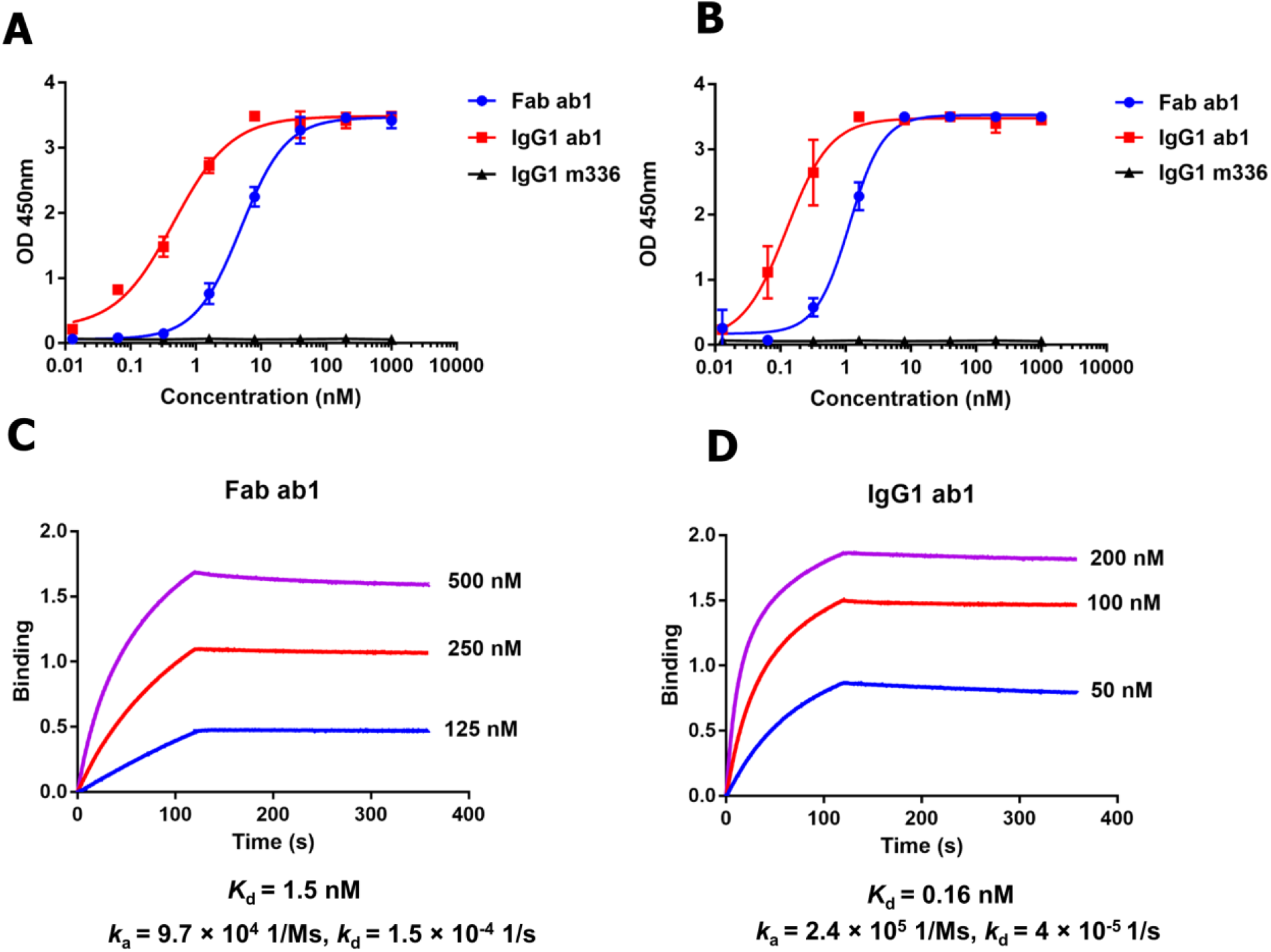
Binding of Fab and IgG1 ab1 to SARS-CoV-2 RBD and S1 proteins as measured by ELISA (A and B) and Blitz (C and D). **(A)** Fab and IgG1 ab1 binding to recombinant RBD measured by ELISA. **(B)** Fab and IgG1 ab1 binding to recombinant S1 measured by ELISA; the MERS-CoV antibody IgG1 m336 was used as a negative control. 100 ng of antigen was coated and serially diluted antibody was added after blocking. After washing, the binding was detected by HRP conjugated anti-FLAG (M2 clone) for Fab ab1, and by HRP conjugated anti human Fc for IgG1 ab1. Experiments were performed in duplicate and the error bars denote ± SD, n =2. **(C)** Blitz sensorgrams for Fab ab1 binding to RBD-Fc. RBD-Fc was coated on the protein A sensors and different concentrations of the Fab ab1 were used to bind to the sensors followed by dissociation. The *k*_a_ and *k*_d_ values were obtained by fitting based on the 1:1 binding model. **(D)** Sensorgrams for IgG1 ab1 binding to RBD-Fc. Biotinylated RBD-Fc was coated on Biotin sensor and IgG1 ab1 was used for binding. The *k*_a_ and *k*_d_ values were obtained by fitting to achieve the fitting parameter R^2^ > 0.99.

**Figure 2.**
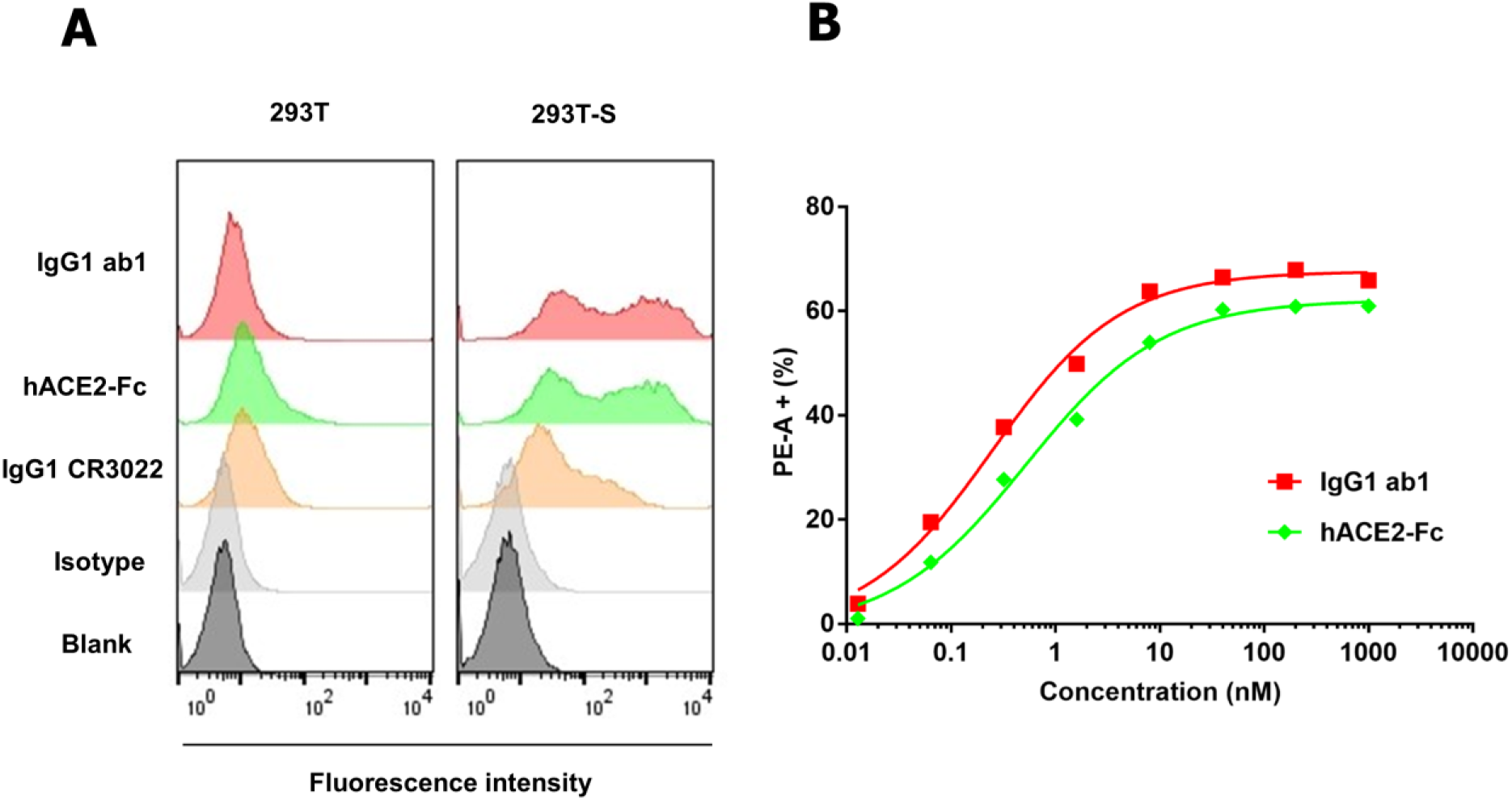
Binding of IgG1 ab1 and hACE2-Fc to 293T cells overexpressing SARS-CoV-2 S (293T-S). (**A**) Binding of IgG1 ab1, hACE2-Fc and IgG1 CR3022 to S transiently transfected 293T cells. The 293T cells without transfection serve as a control. Antibodies or proteins were evaluated at the concentrations of 1 μM. Note the lack of non-specific binding of IgG1 ab1 to 293T cells at this high concentration. (**B**) Concentration-dependent binding of IgG1 Ab1 and hACE2-Fc to 293T-S cells. After 48 h transfection, cells were stained by serially diluted IgG1 ab1 or hACE2-Fc followed by staining using PE conjugated anti human Fc antibody. The percentage of PE-A positive cells is gated and plotted against the antibody concentration. Binding EC_50_ was obtained by using the non-linear mode in Graphpad Prism 7. IgG1 ab1 showed higher binding avidity to 293T-S cells than hACE2-Fc (0.25 nM *v.s.* 0.52 nM for IgG1 ab1 and hACE2-Fc to achieve 50% binding, respectively).

**Figure 3.**
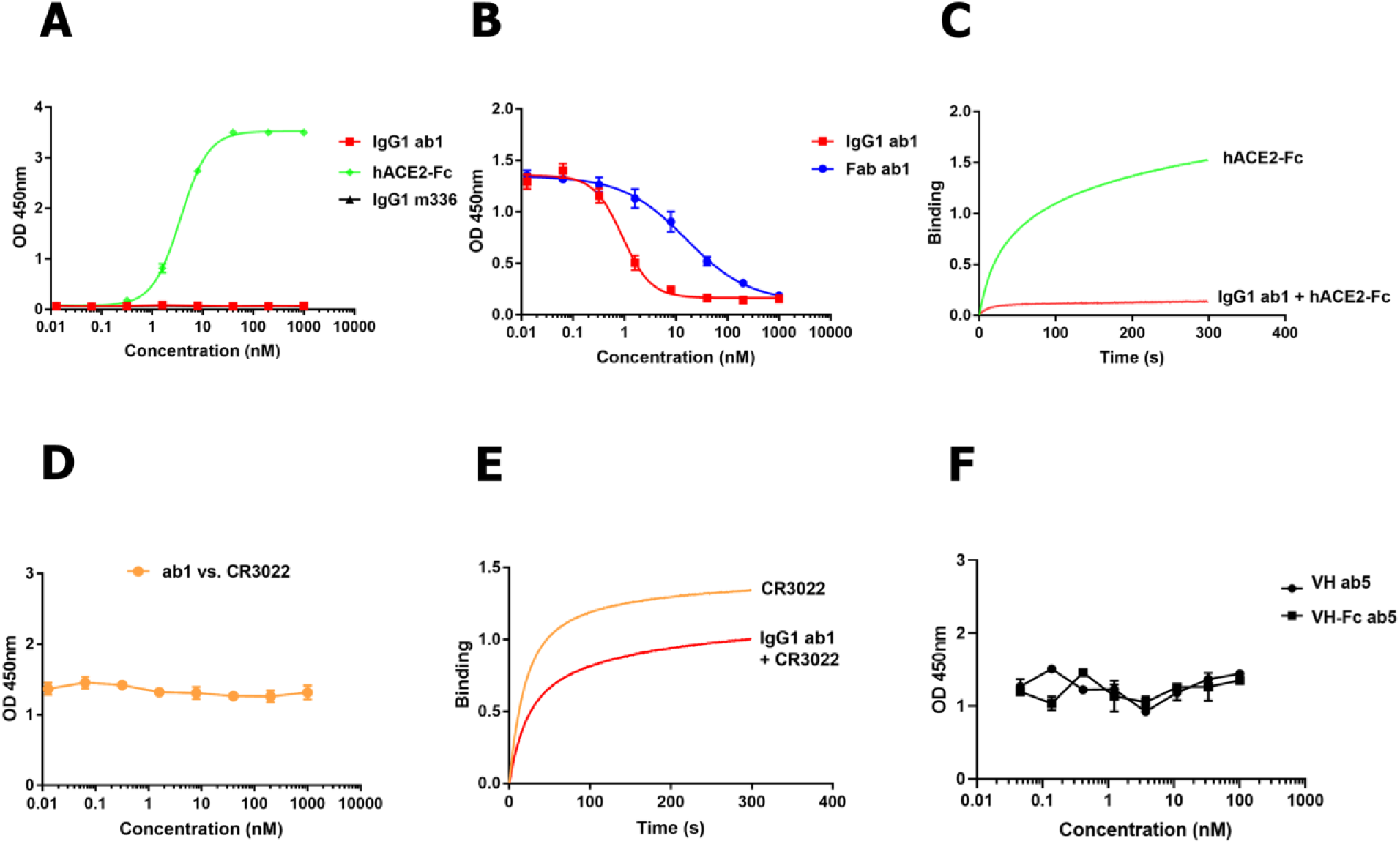
Lack of binding of IgG1 ab1 to SARS-CoV S1 and competition of ab1 with hACE2, CR3022 and ab5 for binding to SARS-CoV-2 RBD as measured by ELISA and biolayer interferometry (Blitz). **(A)** Lack of binding of IgG1 ab1 to SARS-CoV S1 with hACE2-Fc as a positive control and IgG1 m336 as a negative control. **(B)** Competition of ab1 with hACE2 for binding to SARS-CoV-2 RBD; 100 ng of RBD-Fc was coated and 5-fold serially diluted IgG1 or Fab ab1 were added in the presence of 2 nM hACE2-mFc (mouse Fc) followed by PBST washing. For detection, an HRP conjugated anti mouse Fc antibody was used. **(C)** Competition of ab1 with hACE2 tested by Blitz. 100 nM hACE2-Fc was monitored to bind ab1 saturated sensors (red line), which is compared to its independent binding signal to RBD sensor in the absence of ab1(green line). **(D)** Competition ELISA between Fab ab1 and CR3022. ∼10 nM IgG1 CR3022 was incubated with RBD-his in the presence of different concentrations of Fab ab1. After washing, detection was achieved by using HRP conjugated anti human Fc antibody. **(E)** Competition of ab1 with CR3022 tested by Blitz. 100 nM Fab CR3022 was monitored to bind ab1 saturated sensors (red line). The signal was compared to the same concentration of CR3022 binding to the RBD sensor in the absence of ab1 (yellow line). The percentage of signal for CR3022 + ab1 to that of CR3022 alone is ∼77%. Thus, there is no competition between CR3022 and ab1. **(F)** Competition ELISA between ab1 and ab5. ∼6 nM biotinylated IgG1 ab1 was incubated with RBD-Fc in the presence of different concentrations of VH ab5 or VH-Fc ab5. After washing, detection was made by using HRP conjugated streptavidin. Experiments were performed in duplicate and the error bars denote ± SD, n =2.

IgG1 ab1 potently neutralized SARS-CoV-2 pseudovirus with an IC_50_ of 10 ng/ml (Fig 4A). It did not enhance pseudovirus infection of FcγRIA overexpressing 293T-hACE2 cells at any concentration (Fig 4B). It also did not mediate pseudovirus infection of FcγRII expressing K562 cells (Fig S4B). Importantly, IgG1 ab1 exhibited potent neutralizing activity against authentic SARS-CoV-2 in two independent assays -a microneutralization-based assay (100% neutralization at < 400 ng/ml) (Fig. 4C) and a luciferase reporter gene assay (IC_50_ = 200 ng/ml) (Fig. 4D). In agreement with the specificity of binding to the SARS-CoV-2 S1 and not to the SARS-CoV S1 the IgG1 ab1 did not neutralize live SARS-CoV (Fig. 4C). The IgG1 m336 (5) control which is a potent neutralizer of MERS-CoV, did not exhibit any neutralizing activity against SARS-CoV-2 (Fig. 4C). The VH ab5 and VH-Fc ab5 bound the RBD with high affinity and avidity (Fig. S5A.B) but did not compete with hACE2 (Fig. S5C) or neutralize SARS-CoV-2 (Fig. 4D), indicating that not all antibodies targeting epitopes on the RBD affect virus replication.

**Figure 4.**
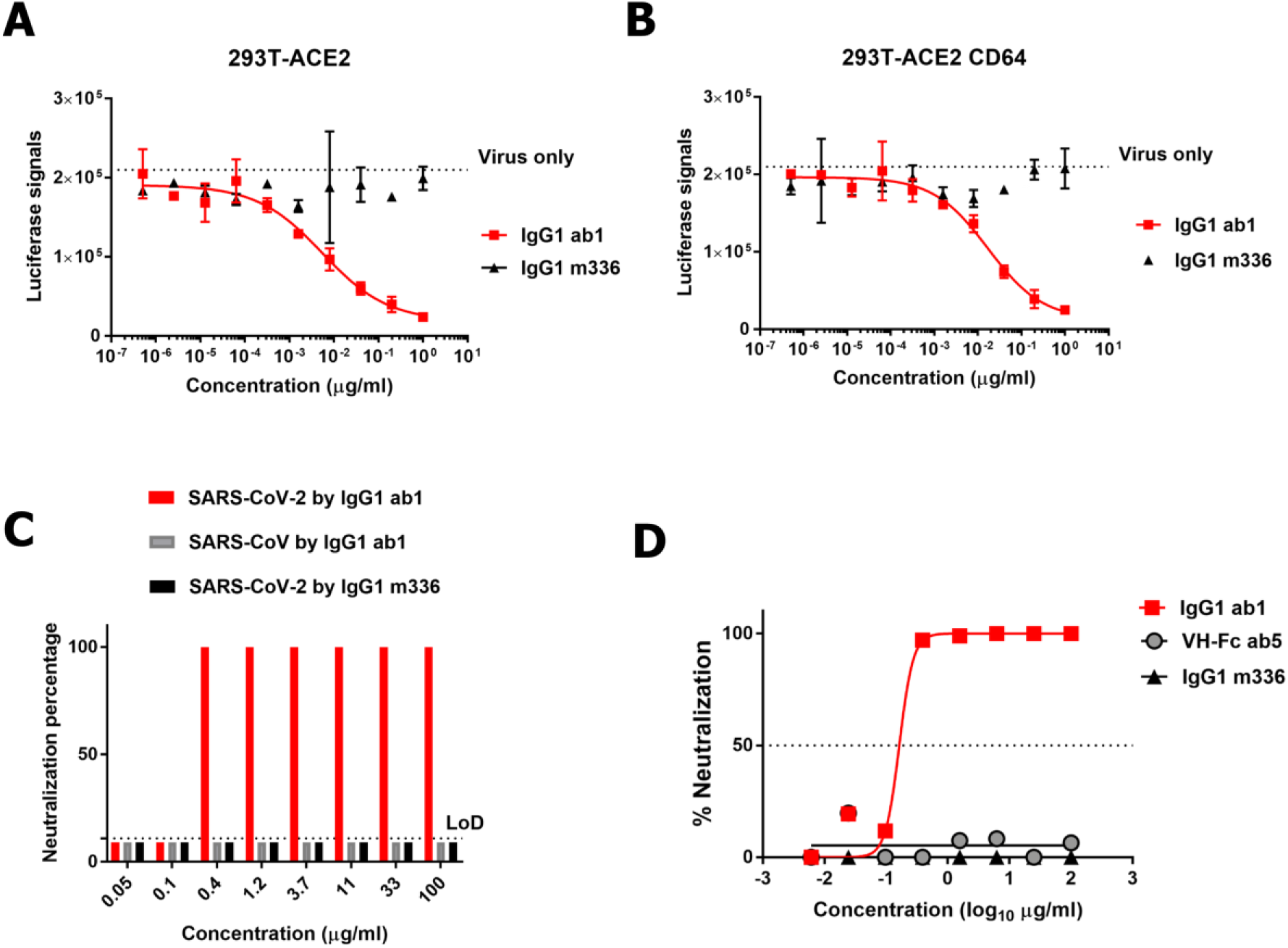
Neutralization activity of IgG1 ab1 and VH-Fc ab5 against SARS-CoV-2 pseudovirus and live virus measured by two different assays. **(A)** Neutralization of SARS-CoV-2 pseudovirus by IgG1 ab1. Lentivirus pseudotyped with SARS-CoV-2 S was produced with luciferase gene. The virus entry into 293T-hACE2 cells was monitored by luminescent signals. (B) Evaluation of ADE for IgG1 ab1 using FcγRIA (CD64) overexpressing 293T-hACE2 cells. The stable cell line 293T-ACE2 transiently expressing CD64 was used for evaluation of pseudovirus entry in the presence of IgG1 ab1 at different concentrations (lowest concentration of 5 × 10^−7^ μg/ml). **(C)** Neutralization of live virus by a microneutralization assay. IgG1 ab1 was pre-mixed with live virus before adding to the VeroE6 cells. After incubation for 4 days at 37°C, virus-induced cytopathic effects were observed under the microcopy. The neutralization capacity was expressed as the lowest concentration capable of completely preventing virus induced cytopathic effect in 100% of the wells. The neutralization of SARS-CoV live virus by IgG1 ab1 was also evaluated. The MERS-CoV antibody m336 was used as a negative control. **(D)** Neutralization of live SARS-CoV-2 by a reporter gene assay. Full-length viruses expressing luciferase were designed and recovered via reverse genetics. IgG1 ab1 was incubated with SARS-UrbaninLuc, and SARS2-SeattlenLuc viruses and then mixed with Vero E6 USAMRID cells in duplicate. Following 48 hours infection, cells were lysed and luciferase activity was measured via Nano-Glo Luciferase Assay System (Promega). Neutralization IC_50_ were defined as the sample concentration at which a 50% reduction in RLU was observed relative to the average of the virus control wells.

To evaluate the efficacy of IgG1 ab1 in vivo we used two animal models. The first one is based on the recently developed mouse ACE2 adapted SARS-CoV-2 which has two mutations Q498T/P499Y at the ACE2 binding interface on RBD (20). IgG1 ab1 protected mice from high titer intranasal SARS-CoV-2 challenge (10^5^ pfu) of BALB/c mice in a dose dependent manner (Fig 5A). There was complete neutralization of infectious virus at the highest dose of 0.9 mg, and statistically significant reduction by 100-fold at 0.2 mg; there was a trend for reduction at 0.05 mg dose but did not reach statistical significance. The IgG1 m336 which potently neutralizes the MERS-CoV in vivo was used as an isotype control because it did not have any activity in vitro. These results also suggest that the RBD double mutations Q498T/P499Y do not affect IgG1 ab1 binding. The second model we used is the transgenic mice expressing human ACE2 (hACE2) (21). Mice were administered 300 ug of IgG1 ab1 prior to wild type SARS-CoV-2 challenge followed by detection of infectious virus in lung tissue 2 days later. Replication competent virus was not detected in four of the five mice which were treated with IgG1 ab1 (Fig 5B). All six control mice and one of the treated mice had more than 10^3^ PFU per lung. These results show clear evidence of a potent preventive effect of IgG1 ab1 in vivo. The reason for absence of virus neutralization in one of the mice is unclear but may be due to individual variation in antibody transfer from the peritoneal cavity where it was administered to the upper and lower respiratory tract. Our previous experiments with transgenic mice expressing human DPP4 and treated with two different doses of m336 (0.1 and 1 mg per mouse) showed similar lack of protection of one (out of four) mice at the lower dose but at the higher dose all four mice were protected (9) similarly to the results obtained with the mouse adapted SARS-CoV-2. The *in vivo* protection also indicates that IgG1 ab1 can achieve neutralizing concentrations in the respiratory tract. This is the first report of in vivo activity of a human monoclonal antibody against SARS-CoV-2 by using two different mouse models. The results also show some similarity between the two models in terms of evaluation of antibody efficacy. In both models about the same dose of antibody (0.2-0.3 mg) reduced about 100-fold the infectious virus in the lungs. This result now suggests that testing of antibody efficacy could be performed at a larger scale than testing with the hACE2 transgenic mice due to the availability of wild type mice. It also shows robust neutralizing activity of IgG1 ab1 in two different models of infection.

**Figure 5.**
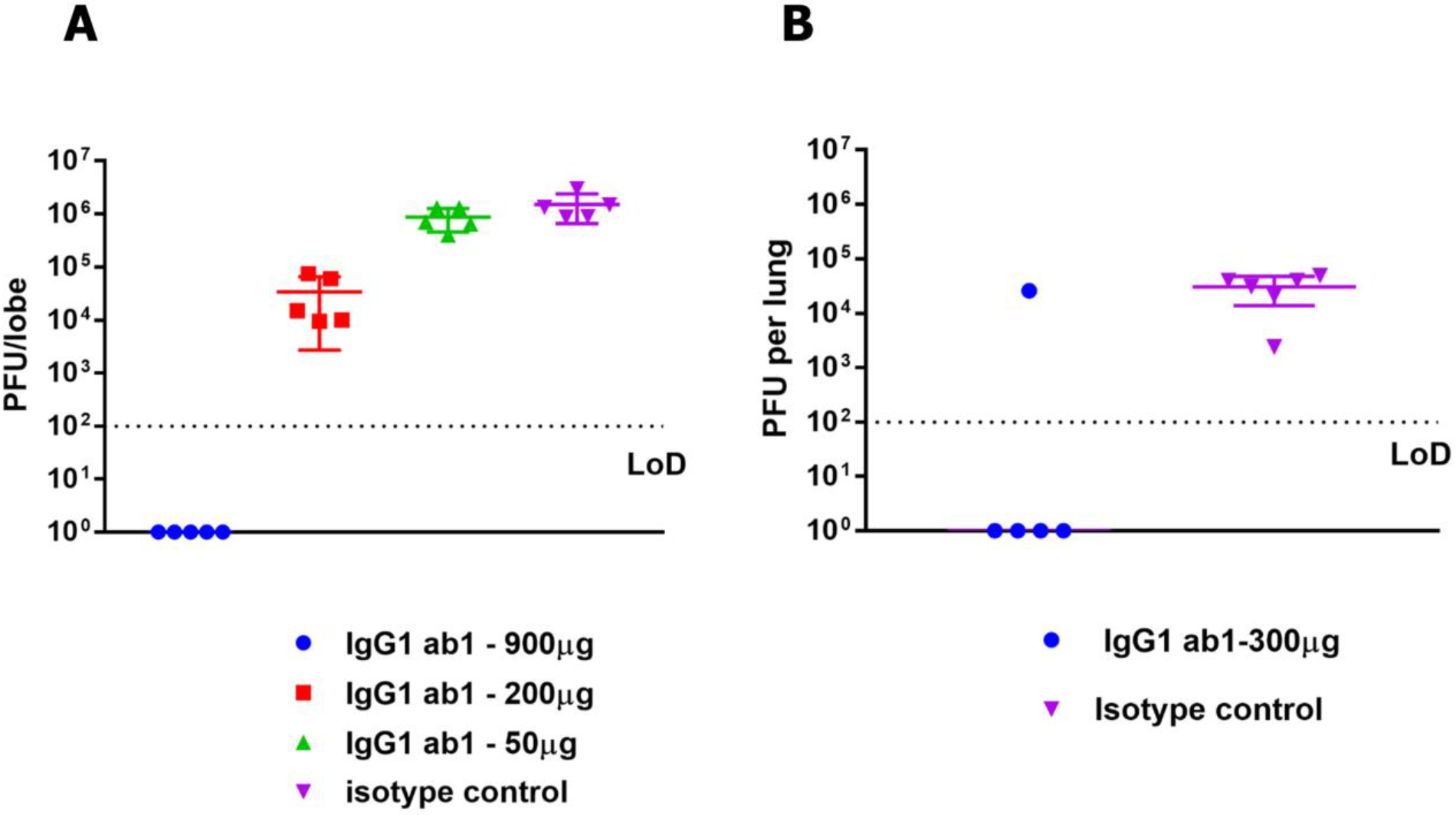
IgG1 ab1 potently neutralizes SARS-CoV-2 in two mouse models. **(A).** IgG1 ab1 inhibits mouse ACE2 adapted SARS-CoV-2 in wild type BALB/c mice. Mice were treated i.p. with varying doses of IgG1 ab1 or an isotype control 12 hours prior to intranasal infection with 10^5^ PFU of mouse adapted SARS-CoV-2. No weight loss was observed over the course of the two-day infection. Lung tissue was homogenized in PBS and virus replication assessed by plaque assay using VeroE6 cells. The assay limit of detection was 100 PFU. **(B)** IgG1 protects hACE2 transgenic mice from SARS-CoV-2 infection. The experimental protocol is similar as the one above except that human ACE2 transgenic mice and wild type SARS-CoV-2 were used. The statistical significance of difference between IgG1 treated and control mice lung infectious virus was analyzed by the one-tailed Mann Whitney *U* test calculated using GraphPad Prism 7.0. A *p* value < 0.05 was considered significant. In the mouse adapted SARS-CoV-2 model, the 0.05 mg treatment group did not show significant differences compared to the isotype controls although minor virus titer decrease was observed. The 0.2 mg and 0.9 mg treatment showed significant decrease compared to the isotype controls (*p*=0.017 and 0.00014, respectively). In the hACE tg mouce model, the *p* value for our comparison was 0.011 and the U value was 2, demonstrating significant difference between IgG1 ab1 treated and control groups.

Interestingly, Fab ab1 had only several somatic mutations compared to the closest germline predecessor genes. This implies that ab1-like antibodies could be elicited relatively quickly by using RBD-based immunogens especially in some individuals with naïve mature B cells expressing the germline predecessors of ab1. This is in contrast to the highly mutated broadly neutralizing HIV-1 antibodies that require long maturation times, are difficult to elicit and their germline predecessors cannot bind native HIV-1 envelope glycoproteins (22, 23). The RBD of the MERS-CoV S protein was previously shown to elicit neutralizing antibodies (24, 25). For SARS-CoV-2 only a few somatic mutations would be sufficient to generate potent neutralizing antibodies against the SARS-CoV-2 RBD which is a major difference from the elicitation of broadly neutralizing antibodies against HIV-1 which requires complex maturation pathways (22, 26–29). The germline-like nature of the newly identified mAb ab1 also suggests that it has excellent developability properties that could accelerate its development for prophylaxis and therapy of SARS-CoV-2 infection (30).

To further assess the developability (drugability) of ab1 its sequence was analyzed online (opig.stats.ox.ac.uk/webapps/sabdab-sabpred/TAP.php); no obvious liabilities were found. In addition, we used dynamic light scattering (DLS) and size exclusion chromatography to evaluate its propensity for aggregation. IgG1 ab1 at a concentration of 2 mg/ml did not aggregate for six days incubation at 37°C as measured by DLS (Fig. 6A); there were no high molecular weight species in freshly prepared IgG1 ab1 also as measured by size exclusion chromatography (SEC) (Fig. 6B). IgG1 ab1 also did not bind to the human cell line 293T (Fig. 2A) even at very high concentration (1 μM) which is about 660-fold higher than its *K*_d_ indicating absence of non-specific binding to many membrane-associated human proteins. The IgG1 ab1 also did not bind to 5,300 human membrane-associated proteins as measured by a membrane proteome array (Fig. 6C).

**Figure 6.**
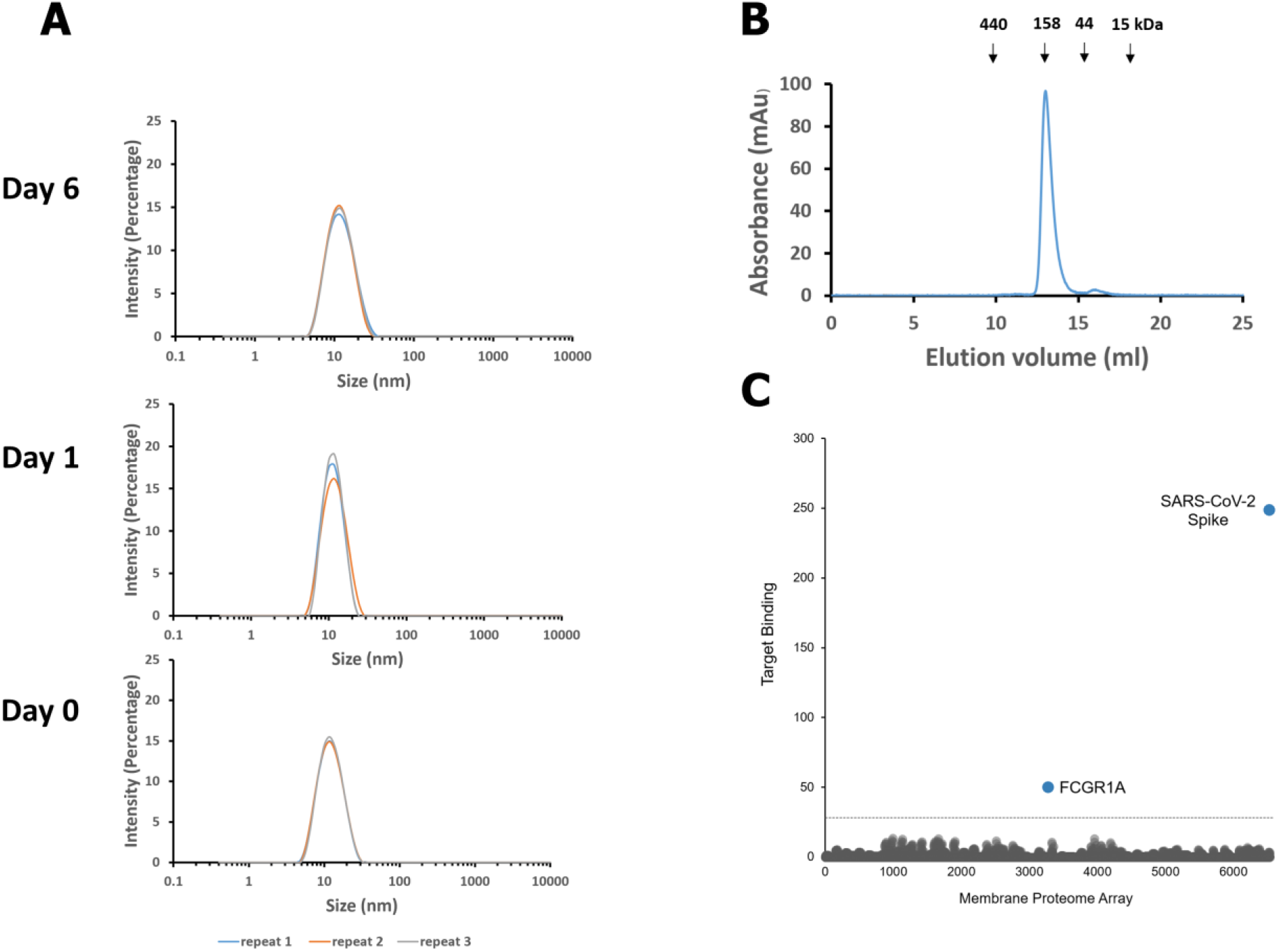
Evaluation of the IgG1 ab1 aggregation by dynamic light scattering (DLS) and size exclusion chromatography (SEC) and test of ab1 non-specificity by a membrane proteome array. **(A)** Evaluation of the aggregation of IgG1 ab1 by DLS. IgG1 ab1 (2 mg/ml) buffered in PBS was incubated at 37°C. On day 0, day 1 and day 6, samples were taken out for DLS measurement to determine the size distribution. All measurements were repeated three times. **(B)** Evaluation of of IgG1 ab1 aggregation by SEC. Size exclusion was performed by loading proteins (150 ul, 1.5 mg/ml) onto the Superdex 200 increase 10/300 GL column. The arrows indicate the peaks of the MW standards in PBS. **(C)** Lack of non-specific binding measured by a membrane proteome array. Specificity testing of IgG1 ab1 (20 μg/ml) was performed using the Membrane Proteome Array (MPA) platform which comprises 5,300 different human membrane proteins, each overexpressed in live cells. To ensure data validity, each array plate contained positive (Fc-binding, FCGR1A; IgG1 ab1 binding, SARS-CoV-2) and negative (empty vector) controls. Identified targets were confirmed in a second flow cytometric experiment by using serial dilutions of the test antibody. The identity of each target was also confirmed by sequencing.

The high affinity/avidity and specificity of IgG1 ab1 along with potent neutralization of virus and good developability properties suggests its potential use for prophylaxis and therapy of SARS-CoV-2 infection. Because it strongly competes with hACE2 indicating a certain degree of mimicry, one can speculate that mutations in the RBD may also lead to inefficient entry into cells and infection. In the unlikely case of mutations that decrease the ab1 binding to RBD but do not affect binding to ACE2 it can be used in combination with other mAbs including those we identified or in bi(multi)specific formats to prevent infection of such SARS-CoV-2 isolates. Ab1 could also be used to select appropriate epitopes for vaccine immunogens and for diagnosis of SARS-CoV-2 infections. The identification of neutralizing mAbs within days of target availability shows the potential value of large antibody libraries for rapid response to emerging viruses.

## Acknowledgments

We would like to thank the members of the Center for Antibody Therapeutics Megan Shi, Cynthia Adams, Du-San Baek and Xioajie Chu for their help with some of the experiments and helpful discussions. We also thank Rui Gong from the Institute of Virology in Wuhan and Rachel Fong from Integral Molecular for helpful suggestions. This work was supported by the University of Pittsburgh Medical Center. David R. Martinez is funded by an NIH NIAID T32 AI007151 and a Burroughs Wellcome Fund Postdoctoral Enrichment Program Award. RSB is supported by grants from the NIH AI132178 and AI108197. Some monoclonal antibodies were generated by the UNC Protein Expression and Purification (PEP) core facility, which is funded by NIH grant P30CA016086.

## Author contributions

DSD, RSB, CTT, JWM and WL conceived and designed the research; WL, ZS and DVZ identified and characterized antibodies; CC made the RBD, ACE2 and other antigens and characterized them; XL and LZ made and characterized reagents and performed some of the experiments; AC performed bioinformatic analysis; EP characterized proteins and helped with the proteome assay; AD and DM performed the neutralization assays; LG, AS and SL performed the animal studies; DSD and WL wrote the first draft of the article, and all authors discussed the results and contributed to the manuscript.

Wei Li, Chuan Chen, Zehua Sun, Doncho Zhelev, John W. Mellors and Dimiter S. Dimitrov are coinventors of a patent, filed by the University of Pittsburgh on March 12 2020, related to antibodies described in this paper. Eric C. Peterson, Alex Conard, John W. Mellors and Dimiter S. Dimitrov are employed by Abound Bio, a company which is developing some of the antibodies for human use.

## Materials and Methods

### Generation, Expression and Characterization of SARS-CoV-2 RBD-Fc, S1-Fc, ACE2-Fc and CR3022 Fab

The SARS-CoV-2 surface glycoprotein and the anti-SARS-CoV antibody IgG1 CR3022 (31) and IgG1 S230 genes were synthesized by IDT (Coralville, Iowa). MERS-CoV-specific IgG1 m336 antibody was expressed in human mammalian cell as described previously (5). The ACE2 gene was ordered from OriGene (Rockville, MD). The RBD domain (residues 330-532) and S1 domain (residues 14-675) and ACE2 (residues 18-740) genes were cloned into plasmid which carries a CMV promotor with an intron, human IgG1 Fc region and Woodchuck posttranscriptional regulatory element (WPRE) to generate the RBD-Fc, S1-Fc and ACE2-Fc expression plasmids. The RBD-avi-his protein with an avi tag followed by a 6×His tag at C-terminal was subcloned similarly. These proteins were expressed with Expi293 expression system (Thermo Fisher Scientific) and purified with protein A resin (GenScript) and by Ni-NTA resin (Thermo Fisher Scientific). The Fab CR3022 antibody gene with a His tag was cloned into pCAT2 plasmid (developed in house) for expression in HB2151 bacteria and purified with Ni-NTA resin. Protein purity was estimated as >95% by SDS-PAGE and protein concentration was measured spectrophotometrically (NanoVue, GE Healthcare).

### Selection, Expression, and Purification of the RBD-specific Fabs and VHs and Conversion to IgG1s or Fc Fusion Proteins

The naïve human antibody phage display libraries were made based on the antibody cDNA from 490 healthy donors peripheral blood monocytes (PBMCs) and splenocytes. The Fab and scFv libraries were constructed by randomly pairing antibody VH and VL gene, and the VH libraries -by grafting CDRs into stable VH scaffolds. These libraries contain very large transformants (size for each ∼10^11^) and are highly diverse (unpublished). For panning, the libraries were preabsorbed on streptavidin-M280-Dynabeads in PBS for 1 h at room temperature (RT) and incubated with 50 nM biotinylated SARS-CoV-2 RBD for 2 h at room temperature with gentle agitation. Phage particles binding to biotinylated antigen were separated from the phage library using streptavidin-M280-Dynabeads and a magnetic separator (Dynal). After washing for 20 times with 1 ml of PBS containing 0.1% Tween-20 and another 20 times with 1 ml of PBS, bound phage particles were eluted from the beads using 100 mM triethanolamine followed by neutralization with 1 M, pH 7.5 Tris-HCl. For the 2^nd^ round of panning, 10 nM (2 nM for the 3^rd^ round) of biotinylated antigen was used as antigen. After the 3^rd^ round of panning against 2 nM biotinylated antigen, 96 individual clones were screened for binding to RBD-Fc fusion protein by phage ELISA. Panels of Fabs and VHs were selected and sequenced. For conversion to Fc-fusion, the VH gene was subcloned into pSecTag B vector (already containing human Fc fragment). For conversion to IgG1, Fab VH and VL gene was inserted into pDR12 vector which contains the IgG1 CH1-CH3 and CL domains. Both VH-Fc and IgG1 were expressed as previously described(32). Protein purity was estimated as >95% by SDS-PAGE and protein concentration was measured spectrophotometrically (NanoVue, GE Healthcare).

### ELISA

The SARS-CoV-2 RBD (residues 330-532) protein was coated on a 96-well plate (Costar) at 100 ng/well in PBS overnight at 4°C. For phage ELISA, phage from each round of panning (polyclonal phage ELISA) or clones randomly picked from the infected TG1 cells (monoclonal phage ELISA) were incubated with immobilized antigen. Bound phage were detected with horseradish peroxidase (HRP) conjugated anti-M13-HRP polyclonal Ab (Pharmacia, Piscataway, NJ). For the soluble Fab/VH binding assay, HRP-conjugated mouse anti-FLAG tag Ab (Sigma-Aldrich) was used to detect Fab/VH binding. For the IgG1 or VH-Fc binding assay, HRP-conjugated goat anti-human IgG Fc (Sigma-Aldrich) was used for detection. For the competition ELISA with hACE2, 2 nM of human ACE2-mouse Fc was incubated with serially diluted Fab, VH, IgG, or VH-Fc, and the mixtures were added to RBD coated wells. After washing, bound ACE2-mouse Fc was detected by HRP-conjugated anti mouse IgG (Fc specific) (Sigma-Aldrich). Another way to do hACE2 competition ELISA is to incubate the fixed concentration of biotinylated hACE2-Fc with gradient concentrations of antibody. After washing, the bound hACE2 was detected by HRP conjugated streptavidin. For the competition ELISA with CR3022, 10 nM IgG1 CR3022 was incubated with serially diluted Fab ab1 or VH ab5, and the mixtures were added to RBD-his coated wells. After washing, bound IgG1 CR3022 was detected by HRP-conjugated anti human Fc antibody. For the competition ELISA between ab1 and ab5, 6 nM biotinylated IgG1 ab1 was incubated with RBD-Fc in the presence of different concentrations of VH ab5 or VH-Fc ab5. After washing, detection was made by using HRP conjugated streptavidin. All the colors were developed by 3,3′,5,5′-tetramethylbenzidine (TMB, Sigma) and stopped by 1 M H_2_SO_4_ followed by recording absorbance at 450 nm. Experiments were performed in duplicate and the error bars denote ± SD, n =2.

### BLItz

Antibody affinities and avidities were analyzed by the biolayer interferometry BLItz (ForteBio, Menlo Park, CA). For affinity measurements, protein A biosensors (ForteBio: 18– 5010) were coated with RBD-Fc for 2 min and incubated in DPBS (pH = 7.4) to establish baselines. 125 nM, 250 nM and 500 nM VH and Fab were used for association. For avidity measurements, RBD-Fc was biotinylated with EZ‐link sulfo‐NHS‐LC‐biotin (Thermo Fisher Scientific, Waltham, MA) (RBD-Fc-Bio). Streptavidin biosensors (ForteBio: 18–5019) were coated with RBD-Fc-Bio for 2 min and incubated in DPBS (pH = 7.4) to establish baselines. 50 nM, 100 nM and 200 nM IgG1 ab1 and VH-Fc ab5 were chosen for association. The association was monitored for 2 min and then the antibody allowed to dissociate in DPBS for 4 min. The *k*_a_ and *k*_d_ were derived from the sensorgrams fitting and used for *K*_d_ calculation. For the competitive Blitz, 500 nM IgG1 ab1 was loaded onto the RBD-Fc coated sensor for 300 s to reach saturation followed by dipping the sensor into the 0.1 μM hACE2-Fc or Fab CR3022 solution in the presence of 500 nM IgG1 ab1. The association was monitored for 300 s. Meanwhile, the signals of 0.1 μM hACE2 or CR3022 binding to the RBD-Fc coated sensor in the absence of IgG1 ab1 was independently recorded. Competition was determined by the percentage of signals in the presence of ab1 to signals in the absence of ab1 (< 0.7 is considered to be competitive) (33).

### Flow Cytometry Analysis

Full-length S protein of SARS-CoV-2 with native signal peptide replaced by the CD5 signal peptide were codon-optimized and synthesized by IDT. S gene was subcloned into our in-house mammalian cell expression plasmid, which were used to transiently transfect 293T cells cultured in Dulbecco’s Modified Eagle’s Medium (DMEM) with 10% FBS, 1% P/S. The transient expression level is tested by FACS staining using the recombinant hACE2-Fc, IgG1 CR3022 and IgG1 ab1. For the determination of binding avidity of IgG1 ab1 and hACE2-Fc to the cell surface S, quantitative FACS was performed by using the 48 h post transfection cells based on standard procedures. Briefly, 5 folds serially diluted antibodies or hACE2-Fc with highest concentration of 1 μM were added into 1×10^6^ cells and incubate at 4 °C for 30 min followed by 3 times washing using PBS + 0.5% BSA buffer (PBSA buffer). Then cells were resuspended to 100 μl PBSA buffer followed by addition of 1 μl PE conjugated anti-human Fc antibody (Sigma-Aldrich) and incubate at 4 °C for 30 min. Cells then was washed by PBSA for 3 times and then analyzed by flow cytometry using BD LSR II (San Jose, CA). The gating of PE-A+ population was performed by FlowJo_V10_CL. The concentrations at which IgG1 ab1 or hACE2-Fc achieved 50% PE-A+ cells (EC_50_) was obtained by fitting using the equation of “[agonist] vs. response -- variable slope (four parameter)” in the Graphpad Prism 7 (San Diego, CA).

### Pseudovirus and Antibody Dependent Enhancement (ADE) of Infection Assays

The ADE assay was based on the SARS-CoV-2 pseudovirus entry into hACE2 expressing cells assay according to previous protocols (34). Briefly, HIV-1 backbone based pseudovirus was packaged in 293T cells by co-transfecting with plasmid encoding SARS-CoV-2 S protein and plasmid encoding luciferase expressing HIV-1 genome (pNL4-3.luc.RE) using polyethylenimine (PEI). Pseudovirus-containing supernatants were collected 48 h later and concentrated using Lenti-X™ concentrator kit (Takara, CA). Pseudovirus neutralization assay was then performed by incubation of SARS-CoV-2 pseudovirus with 5 folds serially diluted antibody with starting concentration of 1 μg/ml for 1 h at 37 °C, followed by addition of the mixture into pre-seeded hACE2 expressing cells in duplicate. The mixture was then centrifuged at 1000 × g for 1 hour at room temperature. The medium was replaced 4 hrs later. After 24 h, luciferase expression was determined by Bright-Glo kits (Promega, Madison, WI) using BioTek synergy multi-mode reader (Winooski, VT). The 50% pseudovirus neutralizing antibody titer (IC_50_) was calculated by non-linear fitting the plots of luciferase signals against antibody concentrations in the Graphpad Prism 7. Experiments were performed in duplicate and the error bars denote ± SD, n =2. ADE was determined by comparing luciferase signals of antibody treated groups to those of virus only group. FcγRs expression level was FACS tested by using FITC conjugated anti-FcγR antibody (Invitrogen).

### SARS-CoV and SARS-CoV-2 Microneutralization Assay

The standard live virus-based microneutralization (MN) assay was used as previously described (9, 35–37). Briefly, serially three-fold and duplicate dilutions of individual monoclonal antibodies (mAbs) were prepared in 96-well microtiter plates with a final volume of 60 μl per well before adding 120 infectious units of SARS-CoV or SARS-CoV-2 in 60 μl to individual wells. The plates were mixed well and cultured at room temperature for 2 h before transferring 100 μl of the antibody-virus mixtures into designated wells of confluent Vero E6 cells grown in 96-well microtiter plates. Vero E6 cells cultured with medium with or without virus were included as positive and negative controls, respectively. Additionally, Vero E6 cells treated with the MERS-CoV RBD-specific neutralizing m336 mAb (5) were included as additional controls. After incubation at 37°C for 4 days, individual wells were observed under the microcopy for the status of virus-induced formation of cytopathic effect. The efficacy of individual mAbs was expressed as the lowest concentration capable of completely preventing virus-induced cytopathic effect in 100% of the wells.

### SARS-CoV and SARS-CoV-2 Reporter Gene Neutralization Assay

Full-length viruses expressing luciferase were designed and recovered via reverse genetics and described previously (38, 39). Viruses were tittered in Vero E6 USAMRID cells to obtain a relative light units (RLU) signal of at least 20× the cell only control background. Vero E6 USAMRID cells were plated at 20,000 cells per well the day prior in clear bottom black walled 96-well plates (Corning 3904). MAb samples were tested at a starting dilution 100 μg/ml, and were serially diluted 4-fold up to eight dilution spots. SARS-CoV-UrbaninLuc, and SARS-CoV-2-SeattlenLuc viruses were diluted in separate biological safety cabinets (BSC) in accordance with UNC safety rules and were mixed with serially diluted antibodies. Antibody-virus complexes were incubated at 37°C with 5% CO_2_ for 1 hour. Following incubation, growth media was removed and virus-antibody dilution complexes were added to the cells in duplicate. Virus-only controls and cell-only controls were included in each neutralization assay plate. Following infection, plates were incubated at 37°C with 5% CO_2_ for 48 hours. After the 48 hours incubation, cells were lysed and luciferase activity was measured via Nano-Glo Luciferase Assay System (Promega) according to the manufacturer specifications. SARS-CoV and SARS-CoV-2 neutralization IC_50_ were defined as the sample concentration at which a 50% reduction in RLU was observed relative to the average of the virus control wells. Experiments were performed in duplicate and IC_50_ was obtained by the fitting of neutralization curves with the “sigmoidal dose-response (variable slope)” equation in Graphpad Prism 7.

### Inhibition of mouse adapted SARS-CoV-2 in wild type mice

A recombinant mouse ACE2 adapted SARS-CoV-2 variant was constructed by introduction of two amino acid changes (Q498T/P499Y) at the ACE2 binding pocket in RBD (20). Virus stocks were grown on Vero E6 cells and viral titer was determined by plaque assay. Groups of 5 each of 10 to 12-month old female BALB/c mice (Envigo, #047) were treated prophylactically (12 hours before infection) intraperitoneally with 900 μg, 200 μg, or 50 μg of IgG1 ab1, respectively. Mice were challenged intranasally with 10^5^ PFU of mouse-adapted SARS-CoV-2. Two days post infection, mice were sacrificed, and lung viral titer was determined by plaque assay.

### Evaluation of IgG1 ab1 Protective Efficacy in a hACE2 Mouse Model of Infection

Human ACE2 transgenic 6-9 week old C3B6 mice were treated intraperitoneally with 0.3 mg of antibody (5 mice) or negative controls (6 mice) 15 hours prior to intranasal infection with 10^5^ PFU of SARS-CoV-2. No weight loss was observed over the course of the two-day infection. Lung tissue was homogenized in PBS and virus replication assessed by plaque assay on VeroE6 cells. The assay limit of detection was 100 PFU.

### Dynamic Light Scattering (DLS)

For evaluation of aggregation propensity, IgG1-ab1 were buffer-changed to DPBS and filtered through a 0.22 μm filter. The concentration was adjusted to 2 mg/mL; ∼500 μL samples were incubated at 37 °C. On day 0, day 1 and day 6, samples were taken out for DLS measurement on Zetasizer Nano ZS ZEN3600 (Malvern Instruments Limited, Westborough, MA) to determine the size distributions of protein particles.

### Size exclusion chromatography (SEC)

The Superdex 200 Increase 10/300 GL chromatography (GE Healthcare, Cat. No. 28990944) was used. The column was calibrated with protein molecular mass standards of Ferritin (Mr 440 000 kDa), Aldolase (Mr 158 000 kDa), Conalbumin (Mr 75 000 kDa), Ovalbumin (Mr 44 000 kDa), Carbonic anhydrase (Mr 29 000 kDa), Ribonuclease A (Mr 13 700 kDa). ∼150 ul filtered proteins (1.5 mg/ml) in PBS were used for analysis. Protein was eluted by DPBS buffer at a flow rate of 0.5 ml/min.

### Computational Analysis of Antibody Sequences

IMGT/V-QUEST tool (40) was used to perform immunogenetic analysis of SARS-CoV-2 RBD-specific mAbs. The unrooted circular phylogram tree of our scFv binders was constructed by using the neighbor joining methods through CLC Genomics Workbench 20.0 (https://digitalinsights.qiagen.com). The RBD sequences were obtained from https://www.ncbi.nlm.nih.gov/genbank/sars-cov-2-seqs/. Liabilities were evaluated online (opig.stats.ox.ac.uk/webapps/sabdab-sabpred/TAP.php) (41).

### Membrane Proteome Array Specificity Testing Assay

Integral Molecular, Inc. (Philadelphia, PA) performed specificity testing of IgG1 ab1 using the Membrane Proteome Array (MPA) platform. The MPA comprises 5,300 different human membrane protein clones, each overexpressed in live cells from expression plasmids that are individually transfected in separate wells of a 384-well plate (42). The entire library of plasmids is arrayed in duplicate in a matrix format and transfected into HEK-293T cells, followed by incubation for 36 h to allow protein expression. Before specificity testing, optimal antibody concentrations for screening were determined by using cells expressing positive (membrane-tethered Protein A) and negative (mock-transfected) binding controls, followed by flow cytometric detection with an Alexa Fluor-conjugated secondary antibody (Jackson ImmunoResearch Laboratories). Based on the assay setup results, ab1 (20 μg/ml) was added to the MPA. Binding across the protein library was measured on an iQue3 (Ann Arbor, MI) using the same fluorescently labeled secondary antibody. To ensure data validity, each array plate contained positive (Fc-binding; SARS-CoV-2 S protein) and negative (empty vector) controls. Identified targets were confirmed in a second flow cytometric experiment by using serial dilutions of the test antibody. The identity of each target was also confirmed by sequencing.

### Ethics statement

Human ACE2 transgenic C3B6 mice (6–9 weeks old) and BALB/c mice (10-12 weeks old) were used for all experiments. The study was carried out in accordance with the recommendations for care and use of animals by the Office of Laboratory Animal Welfare (OLAW), National Institutes of Health and the Institutional Animal Care and Use Committee (IACUC) of University of North Carolina (UNC permit no. A-3410-01).

### Statistical analyses

The statistical significance of difference between IgG1 treated and control mice lung virus titers was analyzed by the one-tailed Mann Whitney *U* test calculated using GraphPad Prism 7.0. A *p* value < 0.05 was considered significant.

## Supplementary Figures and Figure Legends

**Figure S1.**
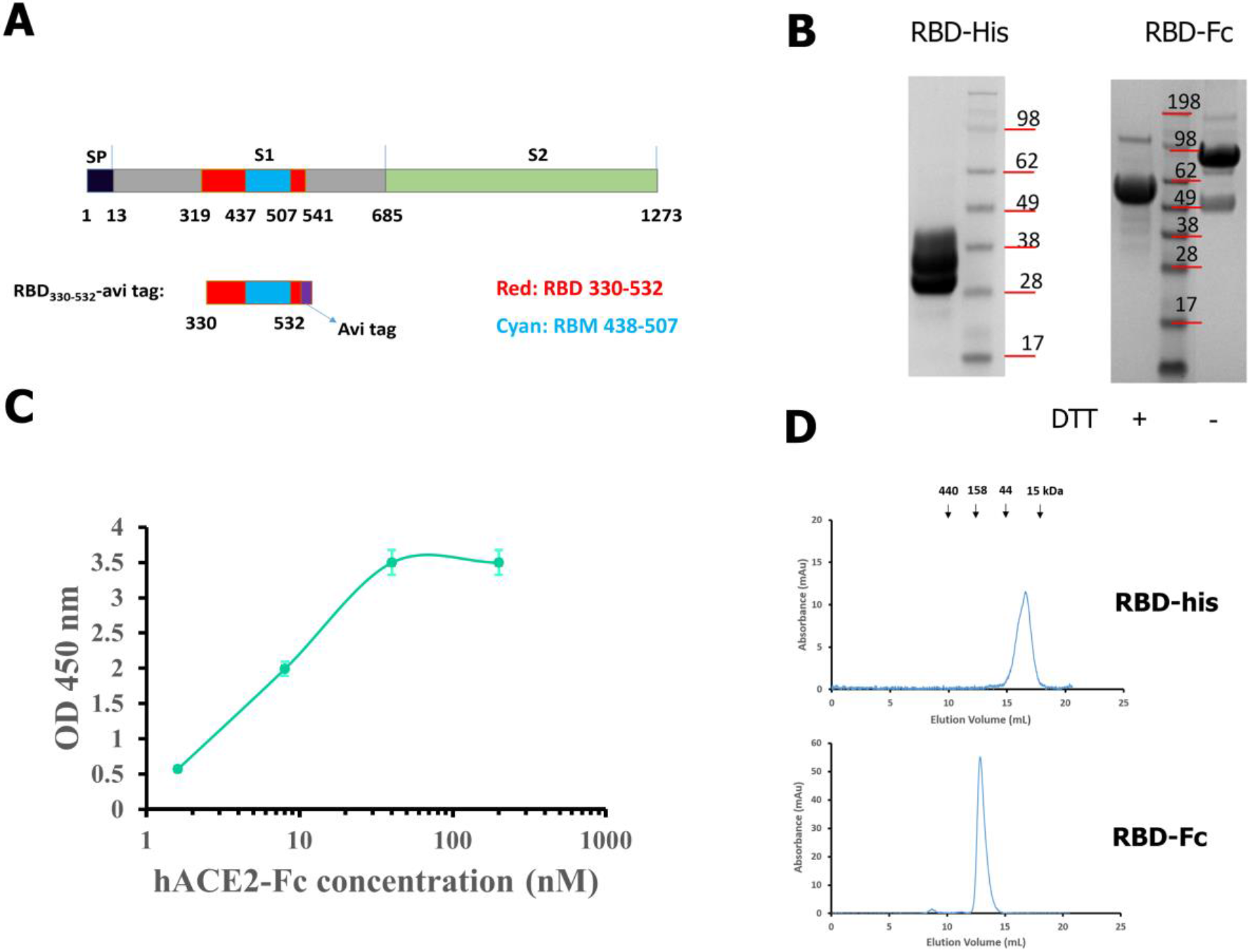
Schematic representation of SARS-CoV-2 S and RBD, and characterization of the RBD as an antigen for panning. **(A)** Schematic representation of the SARS-CoV-2 S and RBD. RBD is highlighted by the red color and the receptor-binding motif (RBM) is pictured by cyan color. RBD_330-532_ is recombinantly expressed in mammalian cells with a C terminal avi tag for in vitro BirA mediated biotinylation. **(B)** SDS-PAGE of RBD-avi-his and RBD-Fc in the presence or absence of DTT. The apparent molecular weight (MW) of RBD-avi-his (heterogeneity ranging from 28 to 38 kDa due to glycosylation) and RBD-Fc (∼100 kDa without DTT and ∼50 kDa with DTT) are consistent with their theoretically calculated MWs. **(C)** ELISA measurement of binding of the recombinant RBD-avi-his to hACE2-mFc (mouse Fc, Sino Biologicals). 50 ng RBD-avi-his was coated on plate with incubation of serially diluted hACE2-mFc. Binding was detected by using HRP conjugated anti-mouse Fc. Experiments were performed in duplicate and the error bars denote ± SD, n =2. **(D)** Evaluation of RBD-his and RBD-Fc by size exclusion chromatography. Size exclusion was performed by the Superdex 200 increase 10/300 GL column. The arrows indicate the peaks of the MW standards in PBS. The well-dispersed single peak indicated RBD-his and RBD-Fc exist as monomers in PBS solution.

**Figure S2.**
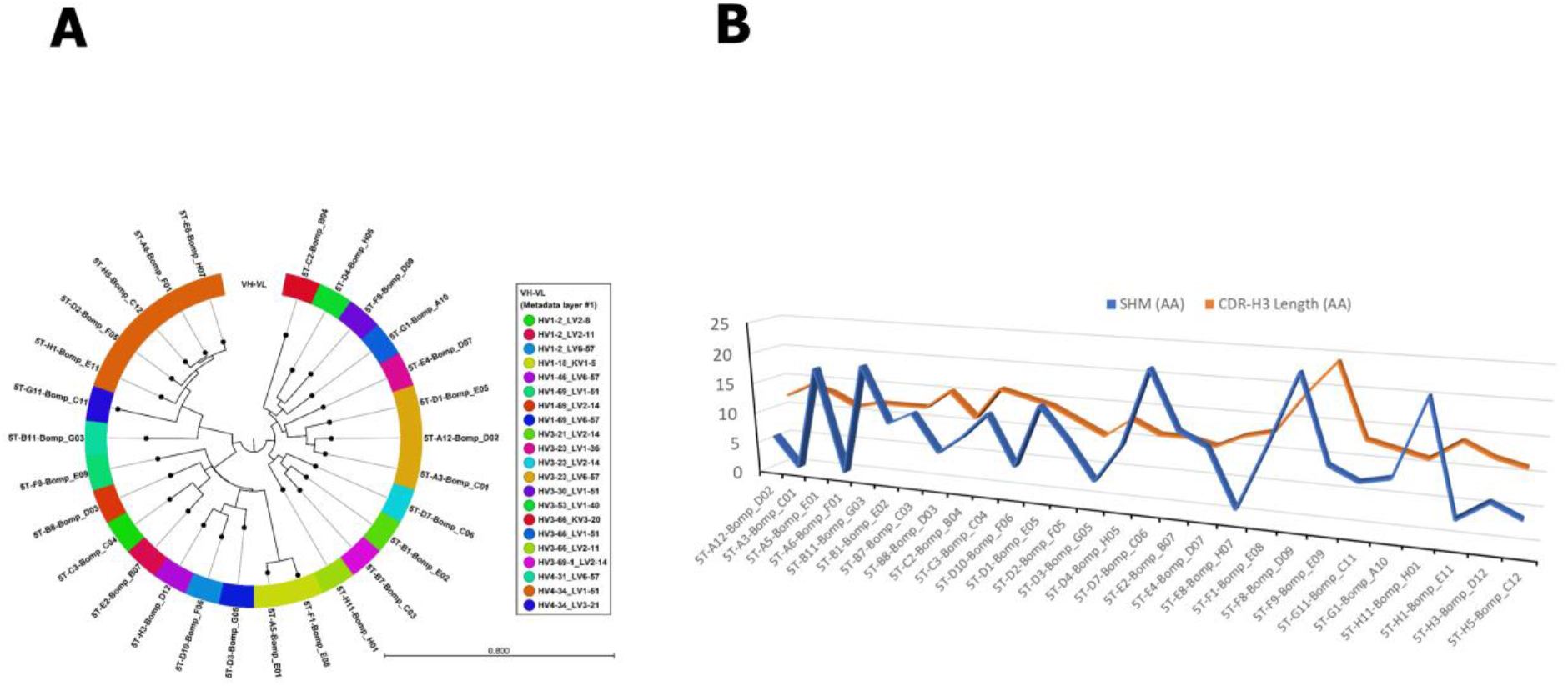
Analysis of the HC/LC usage, and somatic hypermutation (SHM) and CDR-H3 length of 28 SARS-CoV-2 binders selected from our scFv phage library. **(A)** An unrooted circular phylogram tree was constructed using the VH-VL concatenated sequences of 28 unique anti-SARS CoV-2 RBD scFv clones that were mapped, and color coded by IGHV/IGLV germline paring. **(B)** A 3-D line chart showing the number of SHM and CDR-H3 length in amino acid (AA) for each clone.

**Figure S3.**
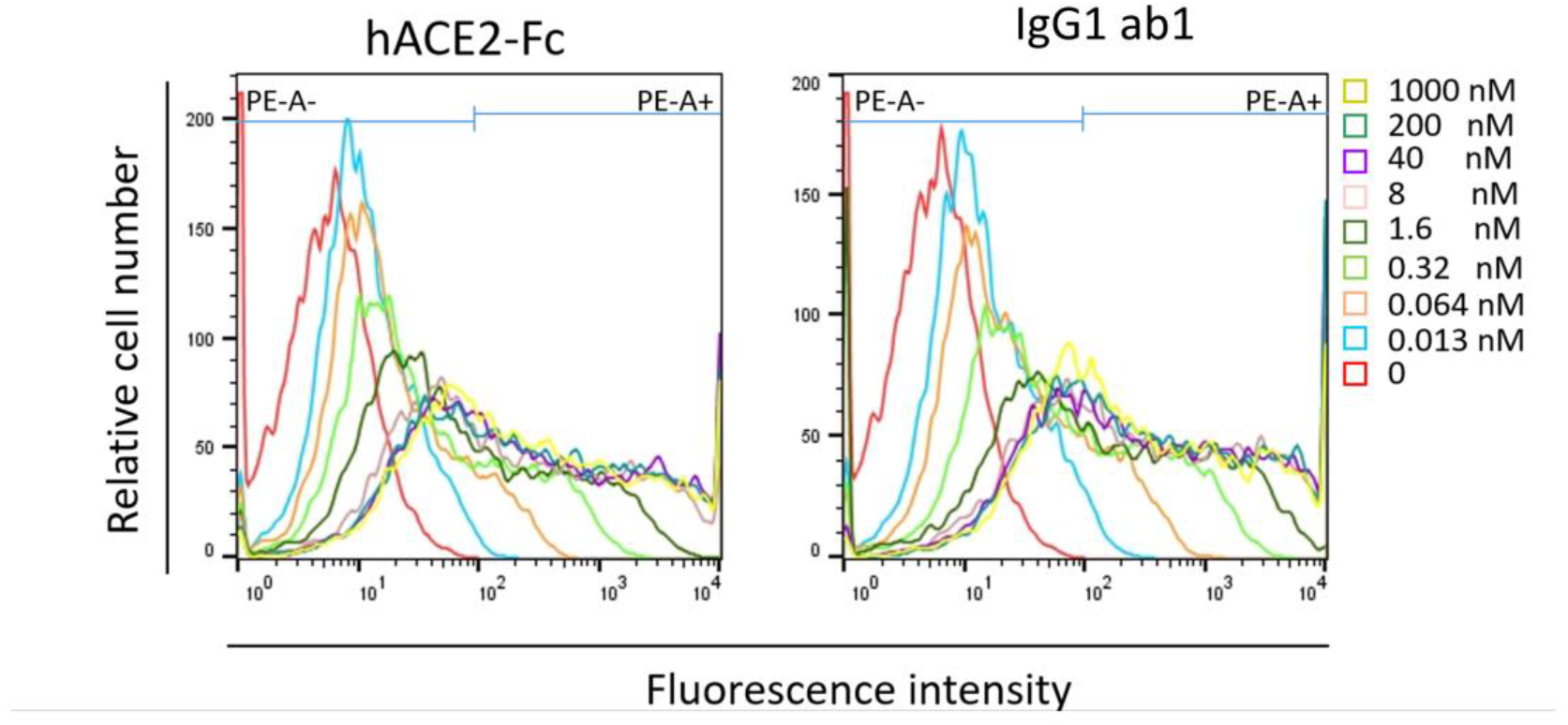
Evaluation of IgG1 ab1 and hACE2-Fc binding avidity to cell surface associated SARS-CoV-2 S by flow cytometry. Cells were incubated with serially diluted antibodies or hACE2-Fc and subsequently with PE conjugated anti-human Fc antibody for flow cytometry analysis. Percentage of PE-A+ cells were defined by the above gate strategy in FlowJ, representing the percentage of IgG1 ab1 and hACE2-Fc bound 293T-S cells.

**Figure S4.**
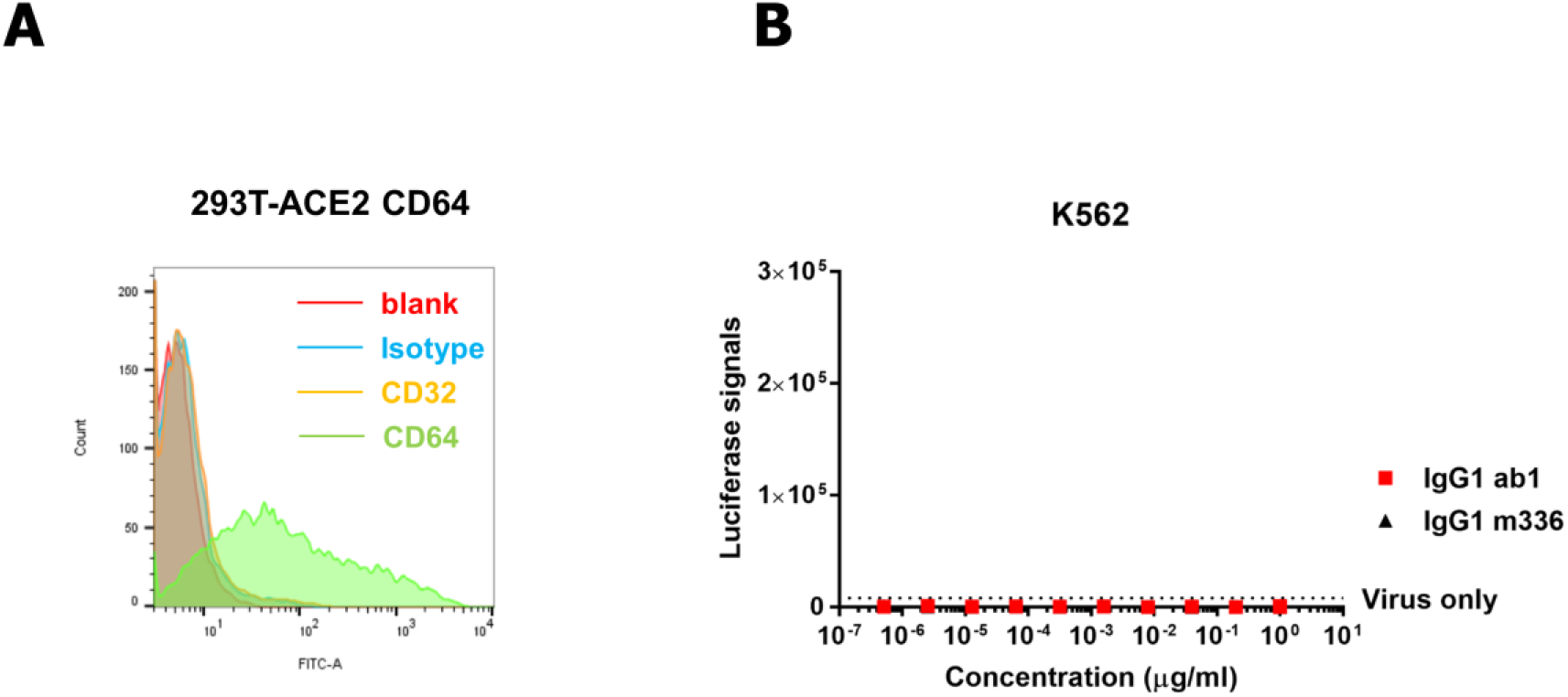
Evaluation of antibody dependentmediated enhancement (ADE) for IgG1 ab1 in FcγR expressing cells. **(A)**293T-ACE2 cells were transiently transfected by FcγRIA, and expression levels after 72 h transfection were detected by FACS by using FITC conjugated anti human CD64 Ab. **(B)** Evaluation of ADE of pseudovirus infection in K562 cells. Pre-mixed varying concentrations of IgG1 ab1 and virus were used to infect K562 cells, which express FcγRII. The infection was monitored by luciferase activity.

**Figure S5.**
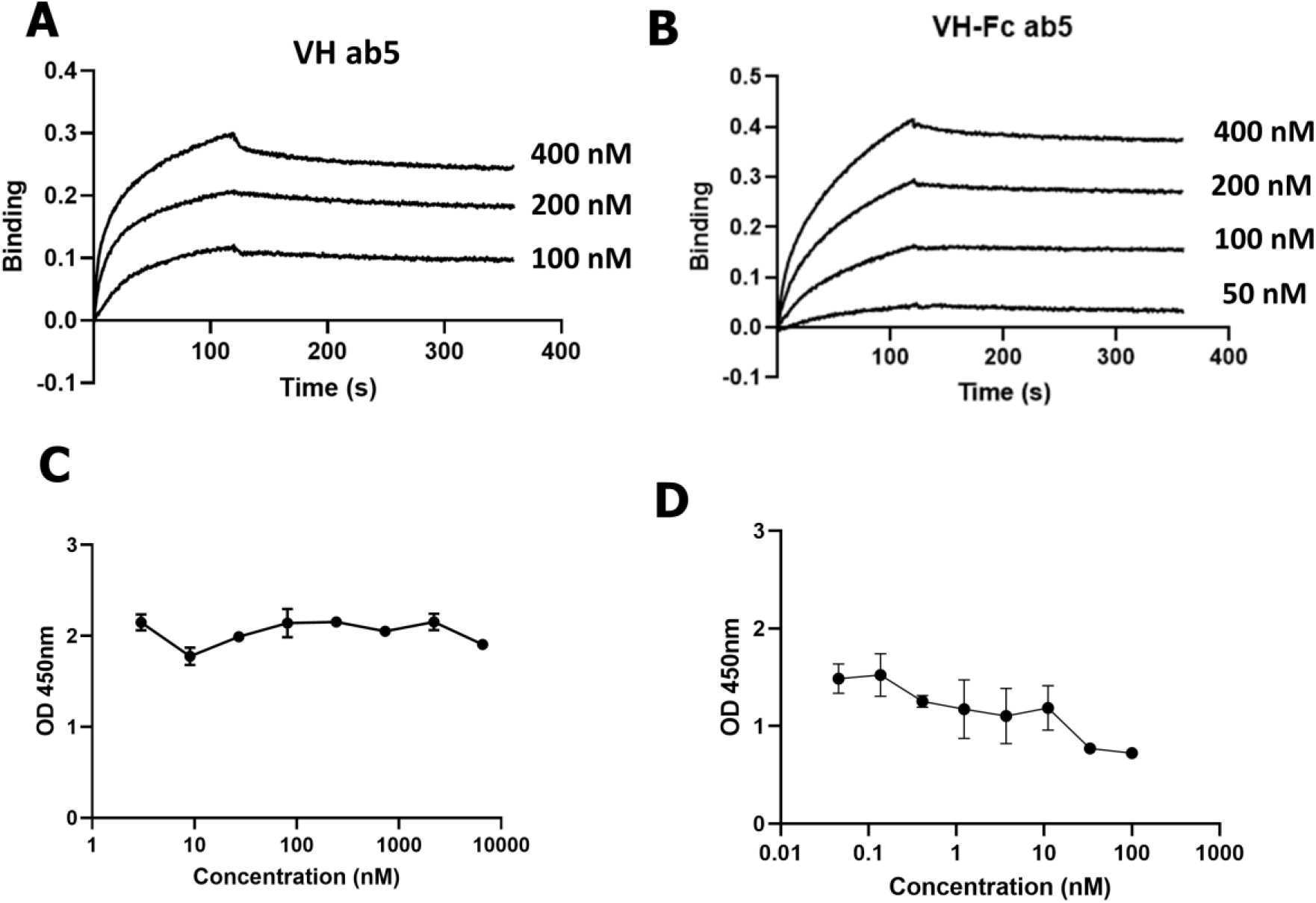
Characterization of VH and VH-Fc ab5 binding to RBD and competition with hACE2 and other antibodies as measured by ELISA and Blitz. **(A)** Blitz sensorgrams for VH ab5 binding to RBD-Fc**. (B)** Blitz sensorgrams for VH-Fc ab5 binding to biotinylated RBD-Fc. Antigens were coated on the protein A or streptavidin sensors and different concentrations of antibody were used to bind to the sensors followed by dissociation. The *k*_a_ and *k*_d_ values were obtained by fitting based on the 1:1 binding model. The equilibrium dissociation constants, *K*_d_, for VH and VH-Fc ab5 were 4.7 nM and 3.0 nM, respectively. **(C)** Competition ELISA between hACE2 and ab5. Biotinylated hACE2-Fc (10 nM) was incubated with RBD-Fc in the presence of different concentrations of VH ab5. After washing, bound hACE2-Fc was detected by using HRP conjugated streptavidin. **(D).** Competition ELISA with CR3022 for binding to SARS-CoV-2 RBD. IgG1 CR3022 (10 nM) was incubated with RBD-his in the presence of different concentrations of VH ab5. After washing, bound CR3022 was detected by using HRP conjugated anti human Fc antibody. Ab5 showed weak competition with CR3022 for binding to SARS-CoV-2 RBD. All the ELISA experiments were performed in duplicate and the error bars denote ± SD, n =2.

